# Ectopic expression of ARGOS8 mediates silk elongation rate response to water deficit in maize (*Zea mays* L.)

**DOI:** 10.64898/2026.07.31.742158

**Authors:** Ignacio R. Hisse, Randy Clark, Jose Rotundo, Andres Reyes, Carla Gho, Jeffrey E. Habben, Mark Cooper, Carlos D. Messina

## Abstract

Water deficit is ubiquitous in maize (*Zea mays* L.) cropping systems worldwide. Ethylene insensitivity in maize has been implicated in improving kernel set and yield under drought, and ARGOS genes modulate ethylene signal transduction by reducing ethylene sensitivity. Given the strong water sensitivity of silk elongation, ARGOS8 overexpression is expected to alter silk growth responses to drought. Experiments were conducted under controlled and field conditions to test the effect of ARGOS8 gene overexpression on silk growth under water deficit. Silk lengths and water use were continuously monitored, and silk elongation rate (SER) response to the fraction of transpirable soil water (FTSW) was evaluated. Silk emergence dynamics were measured in the field by daily counting the silks under contrasting water regimes. ARGOS8 transgenics maintained SER at lower FTSW than controls; however, responses varied among hybrids. Higher SER under water stress translated into a faster silk exertion rate in ARGOS8 transgenics than controls (45 vs. 25 silks d^-1^), leading to a greater number of exerted silks at three days post-silking (424 vs. 377, ∼90% vs. 84% of total silks, *p* < 0.01). Together, these results help clarify the mechanism underlying the ectopically expressed ARGOS8 effect on maize yield improvement under water stress.

**Highlight:** ARGOS8 transgenic expression sustains silk elongation rates and increases silk emergence under water deficit, improving the reproductive performance of maize in water-limited environments.

## Introduction

Reproductive failure in maize is ubiquitous in production environments exposed to stresses around flowering time (Claassen and Shaw, 1970; Cooper *et al*., 2020; Messina *et al*., 2021, 2023) when water deficits are particularly damaging. Developmental and growth processes are implicated in reproductive failure under water deficits, including incomplete gametophyte development (Moss and Downey, 1971), reduced silk elongation (Westgate and Boyer, 1985; Fuad-Hassan *et al*., 2008) and pollination failure (Hall *et al*., 1982), arrest of ovule growth and development (Oury *et al*., 2016b), and kernel abortion (Zinselmeier *et al*., 1995, Zinselmeier *et al*., 1999) due to failure in carbon metabolism and competition (Messina *et al*., 2019). Among physiological traits involved in successful pollination and kernel development, silk elongation rate (SER) integrates critical processes such as silk water potential status, cellular division and expansion, ovary abortion, and hydraulic control mechanisms (Bassetti and Westgate, 1993; Fuad-Hassan *et al*., 2008; Oury *et al*., 2016b; Turc *et al*., 2016). Accordingly, SER constitutes a crucial target for research and maize yield improvement under water deficits.

The single or aggregated effects of abiotic stress on the desynchronization of silk development and emergence have profound effects on kernel set (Messina *et al*., 2019); water use patterns play a pivotal role in the emergence of silks and kernel abortion (Cooper *et al*., 2014a). Presumably, the available empirical evidence points towards a hydraulic effect of the plant nuclear factor Y (NF-Y) on drought adaptation in maize: compared with non-transgenic controls, NF-Y-overexpressing plants maintained higher stomatal conductance and cooler leaf temperatures under water stress (Nelson *et al*., 2007). In the absence of soil moisture data, it is not possible to conclude that a water conservation mechanism was at play. However, improved greenness in pots observed in a dry-down experiment, and enhanced yield under water deficits in field conditions conform well with the hypothesis that improved water status could have favored silk elongation and thus kernel set under drought. Water conservation was implicated in increased maize yields under water deficit conditions likely mediated by an improved water status at flowering time (Cooper *et al.,* 2014a; Messina *et al.,* 2015)

Extended and increased expression of the MADs-box gene *Zmm*28, which begins earlier in maize development and is sustained at higher levels thereafter, enhances photosynthetic capacity, nitrogen uptake, and vegetative growth, resulting in increased grain yield (Wu *et al*., 2019). A related strategy targeted the sugar-signaling metabolite trehalose-6-phosphate (T6P): overexpression of trehalose-6-phosphate phosphatase reduced T6P levels in maize ears, promoting sink strength and increasing growth per ovule, and ultimately increasing yield (Nuccio *et al*., 2015; Zhang *et al*., 2009). Beyond these developmental interventions, hormonal pathways have also been targeted to improve drought tolerance in maize, with the ethylene pathway producing the most consistent results (Guo *et al*., 2014; Habben *et al*., 2014; Simmons *et al*., 2021; Shi *et al*., 2015, 2016). Through ectopic expression of ARGOS (AUXIN-REGULATED GENE INVOLVED IN ORGAN SIZE), genes modulate ethylene signal transduction, reducing the plant sensitivity to ethylene upon overexpression (Shi *et al*., 2015; Rai *et al*., 2015). Overexpression of ARGOS1 has been demonstrated to reduce anthesis-silking interval (ASI) and increase yields, consistent with the negative genetic correlation between ASI response to water deficit and yield (Cooper *et al*., 2014a; Welcker *et al*., 2022) and with the long-term response to selection for yield (Duvick *et al*., 2003; Messina *et al*., 2022).

However, ethylene influences several physiological processes that can directly or indirectly lead to ASI shortening under drought (Hammer *et al*., 2009). For example, ARGOS8 increases maize root elongation at early stages of development (Shi *et al*., 2015, 2017); if this response is maintained for all stages of development and root depths, ARGOS8 overexpression can increase water capture, and thus silk water potential and yield. Ethylene is also implicated in senescence (Young *et al*., 2004), so ARGOS8 could affect leaf area duration and canopy water use, the soil water balance dynamics, and thus the silk water status (e.g., Diepenbrock *et al*., 2022).

Furthermore, ethylene has been implicated in egg maturation via pollination. This event stimulates ethylene production and produce a positive feedback loop to accelerate development (Mol *et al*., 2004). Asynchronous pollination results in asynchronous kernel growth, which increases within-ear kernel competition and induces abortion (Messina *et al*., 2019). Transgenic overexpression of ARGOS8 can reduce ovule sensitivity to ethylene, leading to delayed ovule senescence and, consequently, decreased abortion of ovules and kernels under stress. This mechanism is consistent with the higher kernel numbers per ear (559 vs. 538) observed in transgenic maize hybrids overexpressing ARGOS8 under water deficits (Shi *et al*., 2015).

Because these direct and indirect effects of ethylene, gene × germplasm interactions are expected and have indeed observed (Linares *et al*., 2023, 2024), making genetic engineering for improved drought tolerance in maize a challenging task. Testing the hypothesis that ARGOS8-mediated changes in ethylene sensitivity directly affect silk response to water deficit can guide (i) further inquiry on the regulatory mechanism underpinning silk elongation responses to ethylene and water deficit, and (ii) a better understanding of silking dynamics to improve predictions of gene × germplasm interactions, thereby hastening genetic gain for drought tolerance.

## Materials and methods

### Generation of maize transgenic events

Transgenic maize plants were generated with an *Agrobacterium tumefaciens-*based transformation cassette *ZmGos2-ZmARGOS8*. Expression of *ARGOS8* was controlled by the promoter from the *Zea mays* translation initiation factor *Gos2* gene, *ZmGos2* (de Pater *et al.,* 1992), operationally fused to the intron 1 region from the maize *ubiquitin 1* (*zm-ubi1*) gene (Christensen *et al.,* 1992), conferring moderate constitutive-overexpression. Transcription of the *ARGOS8* gene cassette is completed by the terminator sequence from the *proteinase inhibitor II* (*pinII*) gene from *Solanum tuberosum* (An *et al.,* 1989). Plant transformation was conducted (Unger *et al*., 2001) with two proprietary inbred maize lines (PH184C, PHR03) and resulted in numerous single-copy, backbone-free *ARGOS8* overexpression events. Efficacy of *ZmGos2-ZmARGOS8* transgenic events was previously described in Simmons *et al*. (2021).

### Genetic material

Multiple PH184C and PHR03 events containing the *ZmGos2-ZmARGOS8* construct served as donors for creating water-deficit-tolerant (PH184C hybrids) and a water-deficit-susceptible hybrids (PHR03 hybrids). Single cross proprietary hybrids were created: PHW3G × PH184C, PHW3G ×PHR03, PHEJW × PH184C, and PH11V8 × PH184C and PH1V69 × PH184C. Wild-type (non-transgenic) versions of each hybrid were used as comparators where indicated.

### Experiments and measurements

Two complementary experiments were conducted to evaluate the effects of ARGOS8 transgenes on silk elongation rate and the dynamics of silk emergence under water deficit. A controlled-environment experiment was conducted in Johnston, IA, in a greenhouse-based phenotyping platform to quantify the silk elongation rates. A second experiment was conducted under field conditions in Woodland, CA, to evaluate the dynamics of silk emergence by measuring the exerted silks during the early post-flowering period.

### Experiment 1: Phenotyping platform for silk elongation evaluation

Experiments were conducted in the glasshouse at the Corteva phenotyping platform in Johnston, Iowa, over two years (2014/15 and 2016). Five potted plants per hybrid (one pot per replicate) were grown in 12-L PVC pots filled with a homogeneous soil substrate. Two seeds per pot were sown, and one plant was kept after thinning at three-visible-collar stage (V3; Ritchie *et al*., 1992). Pots were watered daily with a modified one-tenth-strength Hoagland solution containing a complete set of macro- and micronutrients. To minimize soil evaporation, the substrate surface was covered with inert beads, and pots were wrapped with plastic bag.

Air temperature, light, relative humidity, and vapor pressure deficit (VPD) were monitored continuously throughout the experiments. A shielded temperature–humidity probe (Vaisala HMP60) was positioned at canopy height and logged every 5 minutes. Light was monitored with a quantum sensor (LI-190R, LI-COR Biosciences, Lincoln, NE, US) mounted at the height of the uppermost leaves, which recorded instantaneous photosynthetically active radiation (PAR). Supplemental lighting activated automatically when ambient irradiance dropped below a predefined threshold, maintaining PAR levels of 500–700 µmol m^-2^ s^-1^ at midday. Greenhouse setpoints were approximately 24–28°C during the day and 18–22°C at night, with relative humidity typically ranging from 50 to 70%, and VPD kept below 1.5 kPa. All sensors were connected to a Campbell CR1000 datalogger (Campbell Scientific, Logan, UT, USA), and 5-minute readings were averaged to 15-min intervals for consistency with pot-weight measurements described below.

For each pot, soil water content was measured across a gradient from field capacity (i.e., absence of water restrictions) to the lower limit of plant-available water. Pots were kept at field capacity until plants reached silking, after which soil water status was monitored gravimetrically. Pots were placed on individual scales (Ohaus Ranger 3000), and weight was recorded at 15-min intervals during the first week after silking, providing high-resolution estimates of temporal changes in soil water content. To isolate changes attributable to soil water depletion, pot mass measurements were corrected for plant fresh mass using genotype-specific allometric models based on frequently harvested plants. The resulting time series of corrected pot mass was converted to the fraction of transpirable soil water (FTSW) for each pot to quantify the instantaneous soil water status (Koehler *et al*., 2023). For each pot, _FC_ was defined as the pot mass (g) at the onset of the dry-down, when the soil was assumed to be at maximum water-holding capacity after overnight drainage (i.e. field capacity), and _dry_ as the dry reference mass obtained at the end of the dry-down; together, these values defined the transpirable range. FTSW at time was then calculated as:

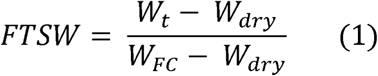

FTSW was constrained to the interval [0,1], with 1 corresponding to field capacity and 0 to the lower limit of plant-available water.

Silking (R1, at least one visible silk in the apical ear) date was recorded from daily observations of individual plants. To avoid the premature arrest of silk elongation caused by ovule fertilization, male inflorescences were removed just prior to the onset of pollen shed (Turc *et al*., 2016). Silk elongation was measured on each plant from one to five days post-silking, using rotational displacement transducers. Each sensor was mounted on a small pulley system carrying a linen thread with a smooth electrical alligator clamp attached at one end to the emerging silks and at the other to a spring to maintain tension. For each plant, at least twenty silks that emerged on the first day of silk emergence were gently aligned and fixed to the thread using the clip. The signal from each transducer was recorded every 15 minutes with a Campbell CR1000 datalogger. Changes in thread displacement over time provided continuous measurements of silk elongation during the post-silking period. Silk elongation rate (SER, mm h^-1^) was calculated as the change in silk length over the elapsed time between consecutive measurements:

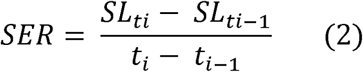

where is the silk length at time *t_i_* and Δ*t = t_i_* -*t_i-1_* is the measurement interval.

### Experiment 2: Field trial for evaluating silk emergence dynamics

A field experiment was conducted in 2016 at a research station in Woodland, California, in a Yolo silt loam to evaluate the silk emergence dynamics of the wild type (WT) and transgenic overexpressing ARGOS8 (ARGOS8) under contrasting water supply conditions. The trials included two irrigation regimes: well-irrigated control and a water-limited treatment, both irrigated via drip lines placed 10 cm below the soil surface. The control was irrigated to meet crop water demand, while the water-limited treatment received approximately 30–40% of crop evapotranspiration from ∼100°Cd pre-silking onwards. The crop was sown on May 1st at a plant density of 8.6 plants m ^2^, with 0.76 m row spacing. Plots were arranged in a randomized complete block design, with three replications. Fertilization, weed control, and pest management were implemented according to local recommendations to minimize confounding stress.

Ten plants per plot were tagged at random prior to silk emergence, and the date of silking was recorded for each tagged plant. The number of exerted silks was then determined daily on these plants from one to six days post-silking. Following Cárcova *et al*. (2000), the exposed portion of the silks was cut from the apical ear of each tagged plant and immediately stored in 90% (v/v) ethanol. Newly exerted silks, identifiable by their bisected apical end, were counted by image analysis (Cooper *et al*., 2014b) to build a cumulative silk-emergence curve for each genotype × water-condition combination.

### Statistical Analysis

Silk elongation responses to decreasing soil water availability were characterized by fitting non-linear mixed models to SER best linear unbiased estimators (BLUEs) as a function of FTSW. A four-parameter logistic function was used to capture the non-linear decline in SER with decreasing FTSW:

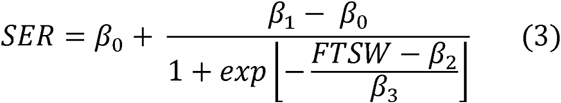

where and (mm h^-1^) represent the lower and upper asymptotes corresponding to SER at very low (FTSW ∼ 0) and near-full (FTSW ∼ 1) soil water availability, respectively; is the inflection point corresponding to the FTSW at which SER changes most rapidly; and describes the steepness of the SER response. Genotype was included as a fixed effect in the model parameters, and replication was modeled as a random effect.

Silk emergence dynamics were analyzed by modeling the cumulative number of exerted silks as a function of time after the first visible silk (1–6 d post-silking). Both water treatments (well-watered and water-limited) were fitted separately, and model selection was based on Akaike Information Criterion (AIC; Akaike, 1974), and biological interpretability. For each water treatment, the silk number on each day was evaluated using linear mixed models with transgene status (i.e. ARGOS8, WT) as the fixed effect, and hybrid and replication as random effects.

Pairwise contrasts were extracted using estimated marginal means, and significance was declared at *p* < 0.05. All mixed-effects models were fitted in R (R Core Team, 2024) using nlme (Pinheiro *et al*., 2024).

## Results

### Wild-type silk elongation response to water deficit is repeatable across years

Continuous monitoring of soil moisture and silk elongation rate made it possible to characterize the SER response to water deficit in detail (Fig. 1). SER of the wild-type hybrid PH1V69 × PH184C was positively related to FTSW, as captured by the fitted sigmoidal function for each year (Fig. 1). The maximum SER estimated by the model, reached as FTSW approached 1, was nearly identical between years (3.25 and 3.30 mm h^-1^ for 2015 and 2016, respectively).

**Fig. 1.**
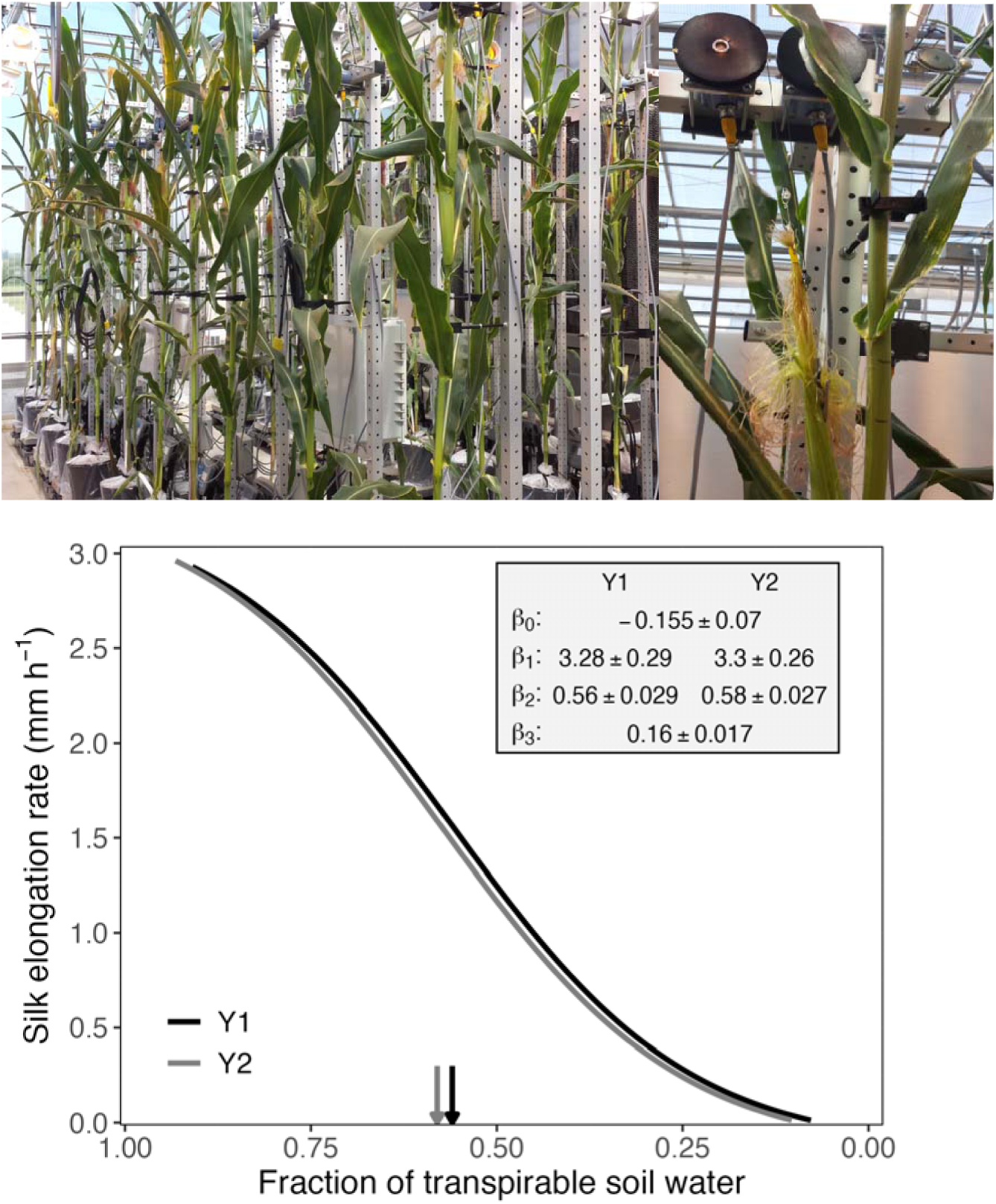
Top: Silk elongation rate phenotyping platform. Data from scales and transducers were recorded using a Campbell CR1000 datalogger. Bottom: Silk elongation rate (SER) response to water deficit for the wild-type hybrid PH1V69 × PH184C, as estimated from the fraction of transpirable soil water (FTSW). Experiments conducted in 2014/15 (Y1) and 2015/16 (Y2) at Johnston, IA. Vertical arrows indicate the FTSW values at which SER responsiveness reaches its maximum, with each arrow corresponding to one experimental year.

The maximum rate of SER increase per unit FTSW, occurring at the inflection point, was 2.78 and 2.71 mm h^-1^ for 2014/15 and 2015/16, respectively, and it was observed at intermediate FTSW (0.56–0.58). The consistency of this response across years demonstrates that the phenotyping approach is highly repeatable.

### Transgenic plants overexpressing ARGOS8 maintain silk elongation rates under severe water deficit, with germplasm-dependent responses

Transgenic plants overexpressing ARGOS8 exhibited higher SER than WT controls across the evaluated range of soil water availability in both the water-deficit-tolerant and -susceptible hybrids (Fig. 2A). Differences between ARGOS8 and WT were more evident in the susceptible background and at low to intermediate soil water levels (FTSW = 0.30–0.60) in both backgrounds (Fig. 2A, C). Specifically, ARGOS8-overexpressing hybrids showed: (i) a steeper SER increase under severe water deficit (FTSW 0–0.25); (ii) a higher maximum SER (Fig. 2A, B), indicating superior silk elongation capacity even under optimal water availability; and (iii) a lower FTSW at the inflection point (Fig. 2A, C), especially for the susceptible hybrid (*p* < 0.05), indicating that the point of maximum SER responsiveness to soil water shifts toward drier conditions. Despite this shift, the SER value at the inflection point remained higher in ARGOS8 than in WT (1.13 vs 1.04 mm h^-1^ in the susceptible hybrid).

**Fig. 2.**
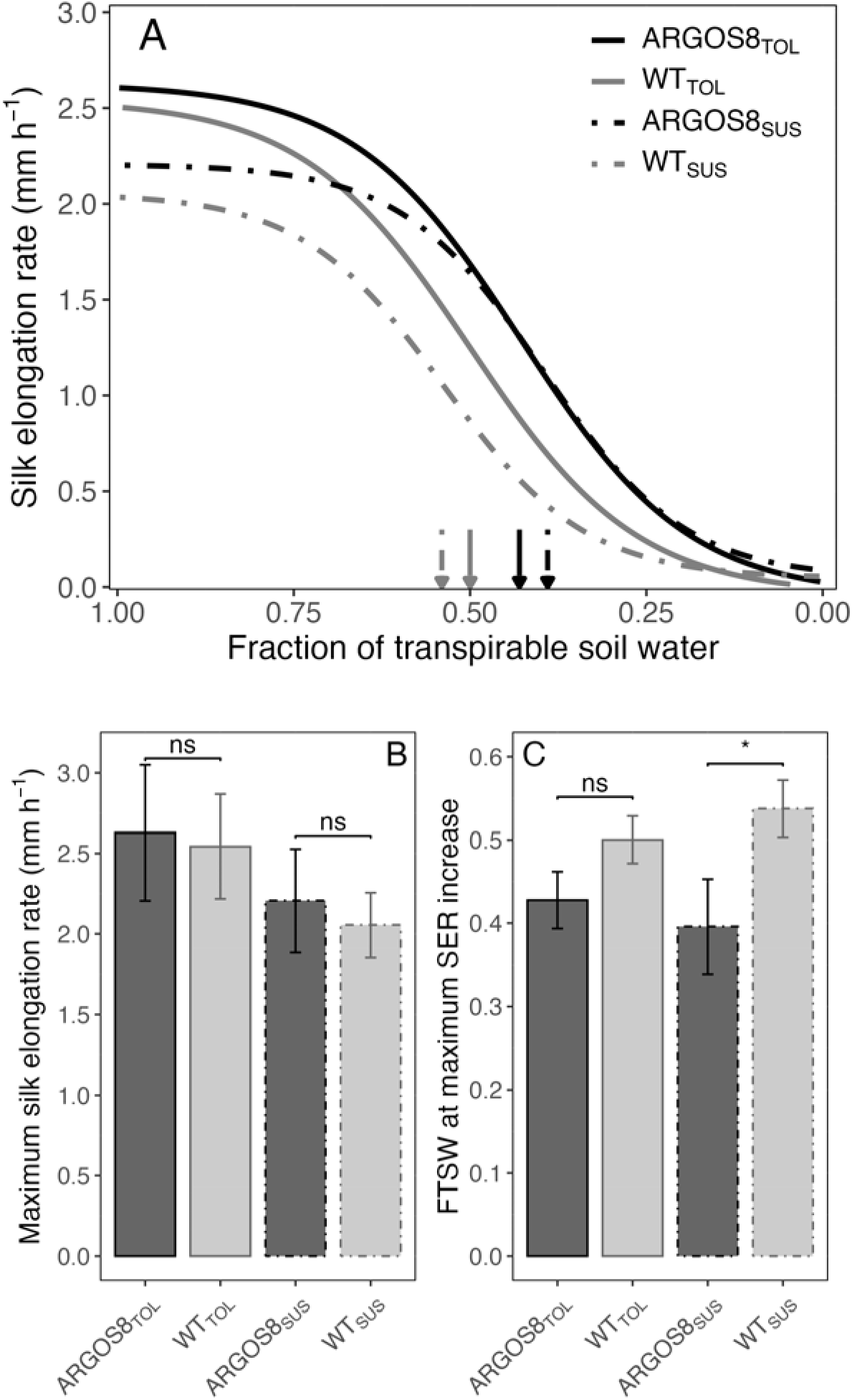
(A) Transgenic plants overexpressing ARGOS8 exhibit a higher silk elongation rate (SER) under water deficit than wild-type (WT) plants. Experiments conducted in 2015/16 included two hybrids: water-deficit-tolerant (PHW3G × PH184C: ARGOS8_TOL_ and WT_TOL_) and water-deficit-susceptible (PHW3G × PHR03: ARGOS8_SUS_ and WT_SUS_), with five random ARGOS8 transformation events for PH184C and four events for PHR03 hybrids. (B) Maximum SER in the absence of water restrictions (parameter β_1_ from A). (C) Fraction of transpirable soil water (FTSW) at the inflection point (parameter β_2_ from A). Bars in B and C represent standard errors. Vertical arrows in (A) indicate the FTSW values at which SER responsiveness reaches its maximum, with each arrow corresponding to each curve. Β_0_: ARGOS8_TOL_ = -0.046, ARGOS8_SUS_ = 0.045, WT_TOL_ = -0.041, WT_SUS_ = 0.045. β_3_: ARGOS8_TOL_ = 0.12, ARGOS8_SUS_ = 0.10, WT_TOL_ = 0.12, WT_SUS_ = 0.10. ns: non-significant; *: *p* < 0.05.

Because the response of ARGOS8-overexpressing plants to water deficit could vary across independent transformation events, the effect of ARGOS8 on SER maintenance was also evaluated for the transformation background PH184C crossed with two testers to produce the hybrids H1 (PH1V69 × PH184C) and H2 (PHEJW × PH184C) (Fig. 3). This result revealed that the effect of ARGOS8 transgenic overexpression upon maintaining SER under water deficit varied between genetic backgrounds introduced by a tester (Fig. 3A). ARGOS8 showed higher SER than WT across the entire range of soil water availability (Fig. 3A), with (i) an increased SER response at low soil water levels (< 0.25 FTSW), (ii) a lower FTSW at the inflection point (Fig. 3A, C), (iii) a higher SER at the inflection point (1.65 vs 1.52 mm h^-1^), and (iv) an increased maximum SER (Fig. 3A, B). The differences disappeared conditioned to the tester since both fitted curves were similar, with comparable maximum SER and FTSW at the inflection point (Fig. 3).

**Fig. 3.**
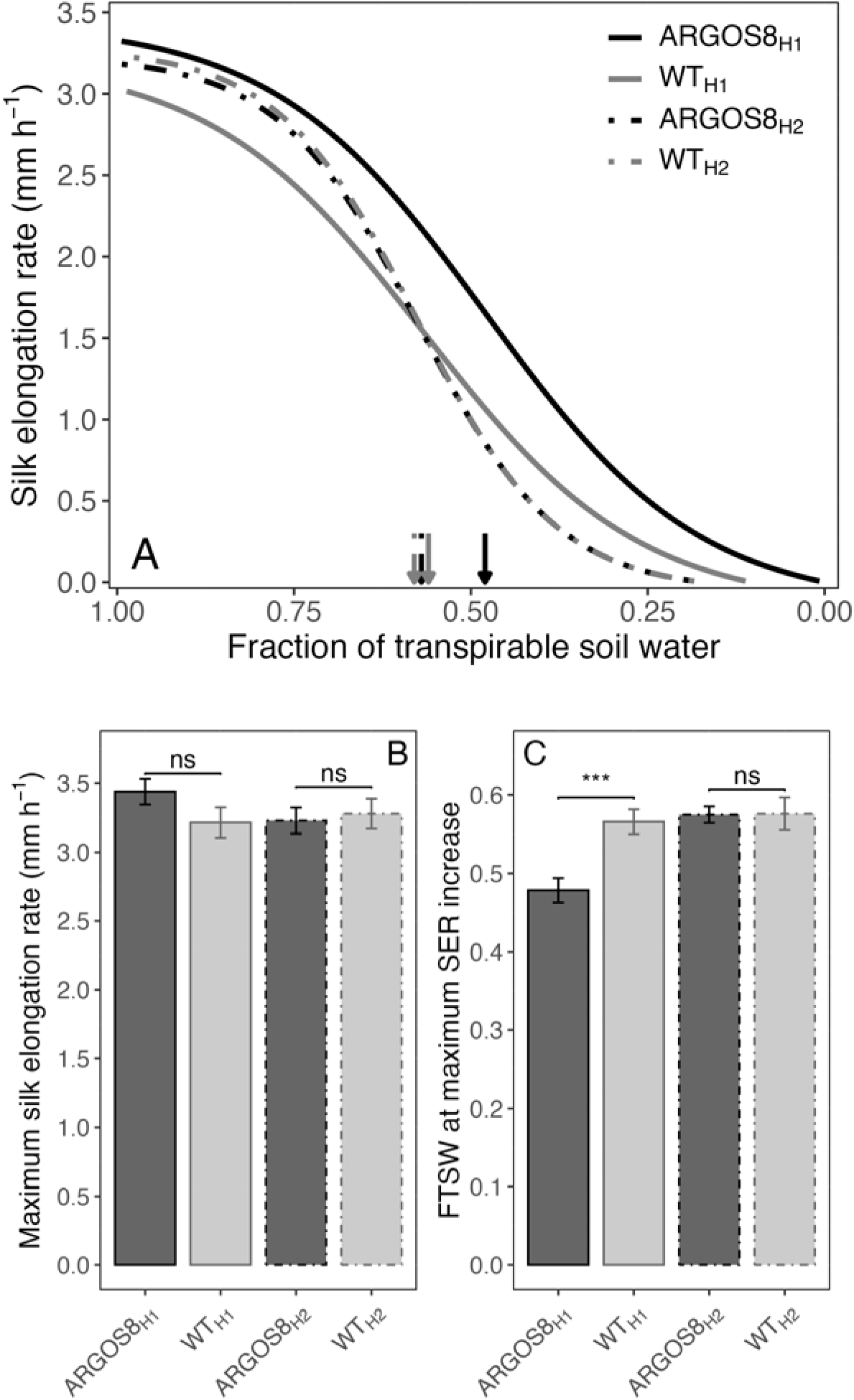
(A) The effect of ARGOS8 transgenic overexpression on silk elongation rate (SER) maintenance under water deficit depends upon the hybrid. Experiments conducted in 2015/16 using five ARGOS8 transformation events for two hybrids: H1 (PHEJW × PH184C) and H2 (PH1V69 × PH184C). (B) Maximum SER in the absence of water restrictions (parameter β_1_ from A). (C) Fraction of transpirable soil water (FTSW) at the inflection point (parameter β_2_ from A). Bars in B and C represent standard errors. Vertical arrows in (A) indicate the FTSW values at which SER responsiveness reaches its maximum, with each arrow corresponding to each curve. β_0_: ARGOS8_H1_ = -0.145, ARGOS8_H2_ = -0.056, WT_H1_ = -0.155, WT_H2_ = -0.054. β_3_: ARGOS8_H1_ = 0.15, ARGOS8_H2_ = 0.10, WT_H1_ = 0.15, WT_H2_ = 0.10. ns: non-significant; ***: *p* < 0.001.

### Ectopic expression of ARGOS8 alters the dynamics of silk emergence under water deficit but not under well-watered conditions

When silk emergence was evaluated from silking to six days post-silking in the tolerant genetic background under water deficit (30–40% of total water demand, Fig. 4B), ARGOS8-overexpressing transgenic plants produced more exerted silks than their respective controls (Fig. 4A). The response patterns also differed between genotypes: ARGOS8 followed a bilinear trajectory with a steeper initial emergence rate (45 silks d^-1^), whereas WT displayed a linear response and lower rate (25 silks d^-1^) (Fig. 4A). Accordingly, significant differences (*p* < 0.05) in emerged silks between ARGOS8 and WT were detected during the early post-flowering period (2–3 d after silking). As a result, by three days post-silking, ARGOS8 had exerted nearly 90% (424 silks) of its total silks, whereas the WT reached 84% (377 silks) (Fig. 4A inset). Compared with the well-watered condition, the decline in the number of emerged silks was more pronounced in WT than in ARGOS8, especially at 2–3 days post-silking (Fig. 4C).

**Fig. 4.**
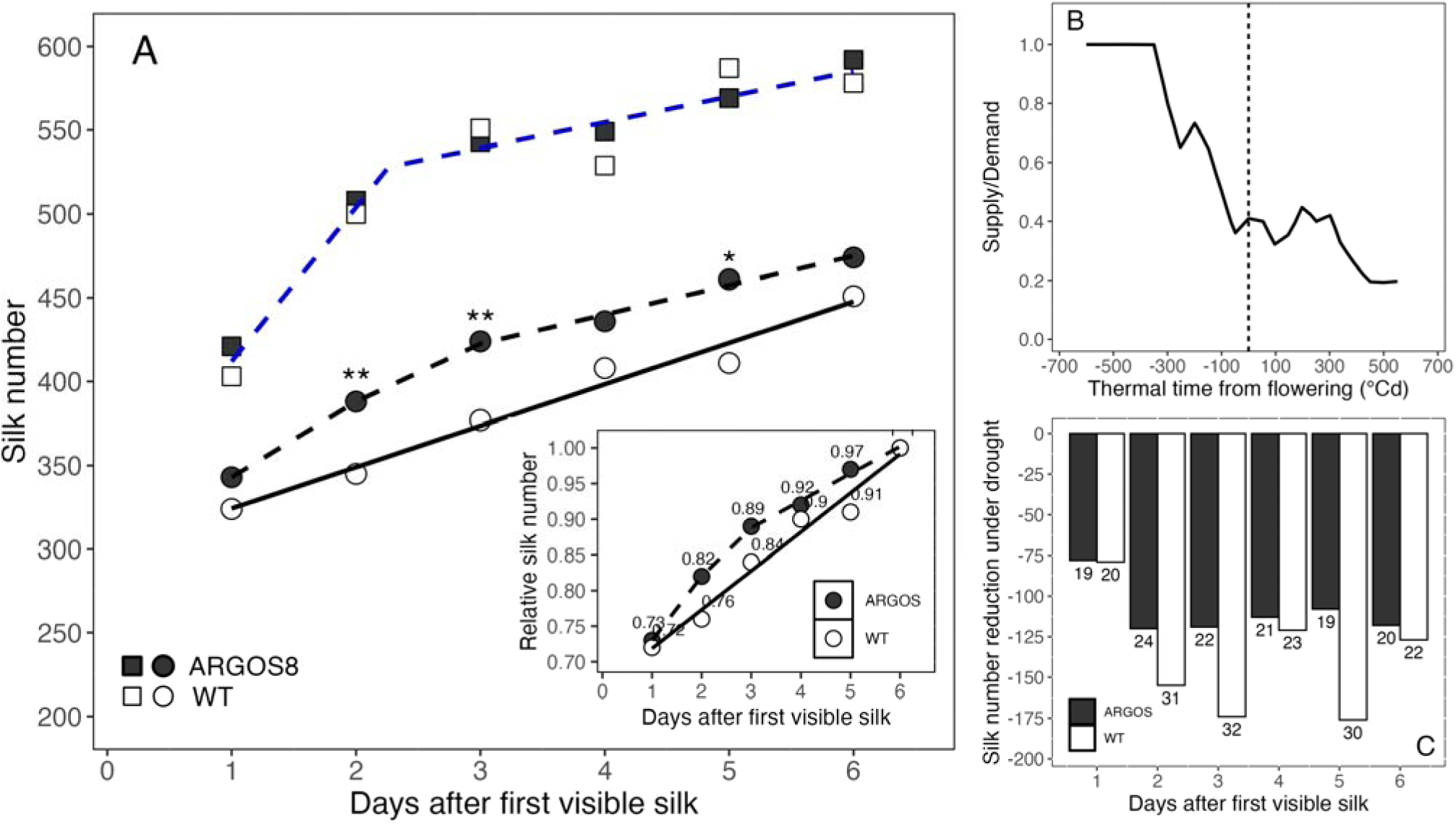
(A) Ectopic expression of ARGOS8 alters the dynamics of silk emergence under water deficit (circles) but not under well-watered (squares) conditions. Experiment conducted in 2016 at Woodland, CA included four ARGOS8 transformation events across four hybrids (PHW3G × PH184C, PHEJW × PH184C, PH1V69 × PH184C, and PH11V8 × PH184C). For the well-watered condition: y = 320 + 92.0x, for x ≤ 2.27; or y = 320 + 92.0(2.27) + 15.3(x−2.27), for x > 2.27. For the water-deficit condition: y_ARGOS8_ = 298 + 45.0x, for x ≤ 2.62; or y_ARGOS8_ = 298 + 45.0(2.62) + 17.5(x−2.62), for x > 2.62. y_WT_ = 299 + 24.7x. Each point represents the mean silk number per plant across hybrids for each genotype (i.e. ARGOS8 and WT) × water condition combination. Asterisks indicate significant genotypic differences at each sampling day (*: *p* < 0.05; **: *p* < 0.01). The inset represents the silk number under water deficit relative to its maximum at day 6 post-silking. (B) Water deficit in the drought-stress treatment, estimated from the water supply to demand ratio. The vertical dashed line indicates silking. (C) Silk number reduction under water deficit relative to well-watered for ARGOS8 and WT across days after the first visible silk. Numbers at the top of each bar indicate the percentage reduction. WT: wild type. Supply to demand ratio (B) is from research station estimates using APSIM (e.g., Cooper et *al.*, 2020; Diepenbrock *et al*., 2022)

## Discussion

In this study, we evaluated the impact of constitutive transgenic overexpression of ARGOS8 on silk response to water deficit by analyzing silk elongation rates and silk emergence dynamics across a wide range of water availability. By phenotyping in both controlled environments and field conditions, which differ in the pattern and rate of water deficit development, we generated direct evidence in both settings rather than relying on extrapolation from controlled-environment results to field conditions (Tardieu and Katerji, 1991; Vadez *et al*. this issue). Our research provides new insights into drought tolerance and the regulation of silk emergence responses to water deficits in maize, paving the way for the development of drought-tolerant maize varieties and improved predictions of transgene × germplasm interactions.

### Transgenic plants overexpressing ARGOS8 maintain silk elongation rates under severe water deficits

Transgenic plants overexpressing ARGOS8 exhibited higher silk elongation rates than WT controls across a broad range of soil water availability, an effect that was more pronounced under low water supply (Fig. 2). Transgenic overexpression of ARGOS family genes has been shown to enhance cell elongation and division, increasing plant and organ size. This results in faster growth rates rather than prolonged growth periods, producing taller plants, larger leaves, and longer ears in maize (Guo *et al*., 2014; Shi *et al*., 2015), as well as larger organs in other species (Feng *et al*., 2011; Hu *et al*., 2003; Kuluev *et al*., 2011). This general growth-promoting effect likely contributes to the higher SER observed in ARGOS8-overexpressing maize even under well-watered conditions (Fig. 2B). The greater SER advantage under low water availability, however, is more consistent with the reduced ethylene sensitivity conferred by ARGOS8 overexpression, a mechanism previously linked to improved drought tolerance (Rai *et al*., 2015; Shi *et al*., 2015). This distinction aligns with previous reports that ARGOS overexpression transgenic events show genotype × environment interactions for maize yield, with the greatest yield benefit typically observed under dry and high temperature conditions (Guo *et al*., 2014; Simmons *et al*., 2021), supporting our findings. This suggests that (i) the environmental interactions previously observed are likely mediated through improved tolerance to drought and related stresses, and (ii) environmental conditions interact with the gene’s effect on complex crop performance traits such as silk elongation and emerged silk number, consistent with the differences observed between ARGOS8 and non-transgenic controls, mostly under water deficit.

### Differences in silk number between ARGOS8 transgenic and WT were observed during the early post-flowering period (2-3 d after silking)

Under water stress, ARGOS8 transgenics exerted more silks than WT controls during the first 3 d after silking (Fig. 4A), consistent with their higher SER, and reflected in the faster silk exertion rate (45 vs. 25 silks d ¹) at this period. Beyond this direct link to SER, silk number is shaped by additional hormonal and physiological factors: ethylene interacts with the stress hormone abscisic acid (ABA) in response to drought, and the development of maize ears and silks is hormonally regulated by ABA concentrations in reproductive tissues (Strader *et al*., 2010; Müller, 2021; Yang *et al*., 2019). Physiological processes, such as (i) turgor pressure and water potential of the silks, which affect silk elongation and development, and (ii) biomass allocation to the ear, largely dependent on plant growth around flowering, also play a role (Bassetti and Westgate, 1993; Andrade *et al*., 1999).

Increased root elongation in ARGOS8 transgenics at early developmental stages (Shi et al., 2015, 2017) enhances water capture, potentially improving silk water potential if this pattern continues at later stages. Additionally, the larger leaves observed in ARGOS1 transgenics (Guo *et al*., 2014) increase light interception, supporting greater plant growth and biomass allocation to the ear, consistent with the shared growth-promoting activity described across the ARGOS gene family. However, carbon limitation resulting from reduced leaf area and photosynthesis, typically affects ovary abortion later in development, around five days after silking (Oury *et al*., 2016a). This is well after the 2–3 d post-silking window in which the increased number of exerted silks was observed in ARGOS8 transgenics. This temporal mismatch suggests that early differences in silk emergence are unlikely to be driven by carbon limitation. Instead, the most plausible explanation is that the enhanced expansive growth of reproductive organs, strongly governed by plant water status, primarily contributed to the greater number of emerged silks in ARGOS8 transgenics during the early post-silking period.

The increased number of silks exerted in ARGOS8 transgenic plants, particularly during early post-flowering period (2-3 days after silking), has significant implications for reproductive success under water stress. First, more silks exerted during the early post-flowering period significantly influence silking dynamics, enhancing the chances of successful pollination (Cárcova *et al*., 2000). This favors more uniform resource allocation among grains by reducing intra-ear competition driven by temporal asynchrony in development among ovary cohorts along the ear (Otegui *et al*., 1995; Cárcova and Otegui, 2007; Messina *et al*., 2019). Second, because the total duration of silk growth, from initiation to cessation, is relatively constant (Fuad-Hassan *et al*., 2008), a faster elongation rate allows silks to clear the husk and become exposed earlier within that fixed growth window, thereby prolonging the period during which they remain exposed and receptive to pollen. This is particularly important under water stress, because the tolerance window, from the appearance of the first silk in a cohort to the point of its abortion, is substantially shortened (Oury *et al*., 2016b; Messina et al., 2019). Consequently, maintaining active silk growth is essential for successful and synchronous fertilization, as it supports pollen tube progression along the silk and reduces the propensity for ovary abortion (Oury *et al*., 2016b).

### The effect of ARGOS8 transgenic overexpression on maintaining silk elongation rate under water deficit was germplasm-dependent

Previous studies have documented substantial natural genetic variation in silk elongation and ASI responses to water deficit (Bolaños and Edmeades, 1996; Cooper *et al.,* 2014a; Turc *et al*., 2016; Messina *et al.,* 2021). This variation is consistent with the extensive differential gene expression in stress signaling, hormonal pathways, and defense metabolism, all of which are critical for adaptation and reproductive success (McNinch *et al*., 2020). Importantly, the estimated effect of a transgene on such quantitative traits is typically influenced by both the genetic background and the environment in which it is expressed (Linares *et al*., 2023, 2024).

Interactions among transgenes, germplasm, and environment have similarly been documented for other agronomic traits, including yield and plant height, under both stress and non-stress conditions (Simmons *et al*., 2021; Linares *et al*., 2023, 2024). This pattern is consistent with the transgene × germplasm interaction detected for SER in our study (Fig. 3). Unlike transgenes developed for insect and weed control, which exhibit relatively simple mechanistic effects and broad effectiveness across diverse environments and genetic backgrounds (Estruch *et al*., 1997; Dill *et al*., 2008; Kumar *et al*., 2008; Duke, 2011), drought tolerance arises from complex physiological processes involving multiple genes and pronounced environmental sensitivity. These pathways often involve more complex interactions that require precision phenotyping, molecular markers to study the effects of multiple genes, and a deep understanding of trait genetic architecture (Cooper *et al*., 2014a; Khaipho-Burch *et al*., 2023).

Despite the consistent effect of ARGOS8 on SER across transformation backgrounds, the germplasm-dependent effects of ARGOS8 transgenes observed here underscore the importance of evaluating transgene performance not only across environments but also within diverse germplasm backgrounds.

## Conclusions

Moderate constitutive overexpression of ARGOS8 enhanced silk growth in maize under water deficit, sustaining higher silk elongation rates and increasing the number of silks exerted during the critical early post-silking period. The higher SER reflects both the growth-promoting activity shared across the ARGOS gene family and the reduced ethylene sensitivity conferred specifically by ARGOS8 overexpression, while the greater number of exerted silks is primarily attributed to enhanced expansive growth of reproductive organs under declining water availability. The magnitude of the transgene effect varied among hybrids, highlighting a clear transgene × germplasm interaction consistent with the complex genetic architecture of drought tolerance. This underscores the need to evaluate drought-tolerance transgene performance across diverse genetic backgrounds and testing conditions. Together, these findings provide mechanistic insight into the role of ethylene-related pathways in silk growth and emergence under water-limited environments.

## Acknowledgements

All experiments were conducted at Corteva Agriscience. We thank Steven Becker and Ross Megargel for their help with engineering and fabrication of the phenotyping platforms, Logan Anderson and Dan Chamberlain for their assistance with experimentation, Yinan Fang for his help with statistical modeling, and Mary Trimnel for her assistance with seed production, John Arbuckle supported the project.

## Author contributions

IRH, RC, MC, and CDM: Conceptualization; AR, CG, RC, and CDM: Data curation; IRH, RC, and CDM: Formal analysis; AR, CG, RC, and JEH: Investigation; RC, JEH, IRH, and CDM: Methodology; CDM, RC, and MC: Project administration; CDM: Supervision; IRH, MC, and CDM: Writing original draft; IRH, JR, JEH, RC, MC, and CDM: Writing review & editing. All authors have read and agreed to the published version of the manuscript.

## Data availability

At the sole discretion of Corteva Agriscience the data could be made available upon request.

## Abbreviations

ARGOS: AUXIN REGULATED GENE INVOLVED IN ORGAN SIZE
ARGOS8: ARGOS family gene 8
FTSW: fraction of transpirable soil water
SER: silk elongation rate
WT: wild type.

